# Blazing Signature Filter: a library for fast pairwise similarity comparisons

**DOI:** 10.1101/162750

**Authors:** Joon-Yong Lee, Grant M. Fujimoto, Ryan Wilson, H. Steven Wiley, Samuel H. Payne

**Author notes:** Corresponding author: (SHP).

## Abstract

Identifying similarities between datasets is a fundamental task in data mining and has become an integral part of modern scientific investigation. Whether the task is to identify co-expressed genes in large-scale expression surveys or to predict combinations of gene knockouts which would elicit a similar phenotype, the underlying computational task is often a multi-dimensional similarity test. As datasets continue to grow, improvements to the efficiency, sensitivity or specificity of such computation will have broad impacts as it allows scientists to more completely explore the wealth of scientific data. A significant practical drawback of large-scale data mining is that the vast majority of pairwise comparisons are unlikely to be relevant, meaning that they do not share a signature of interest. It is therefore essential to efficiently identify these unproductive comparisons as rapidly as possible and exclude them from more time-intensive similarity calculations. The Blazing Signature Filter (BSF) is a highly efficient pairwise similarity algorithm which enables extensive data mining within a reasonable amount of time. The algorithm transforms datasets into binary metrics, allowing it to utilize the computationally efficient bit operators and provide a coarse measure of similarity. As a result, the BSF can scale to high dimensionality and rapidly filter unproductive pairwise comparison. Two bioinformatics applications of the tool are presented to demonstrate the ability to scale to billions of pairwise comparisons and the usefulness of this approach.

## Introduction

Data mining is frequently used in scientific research for hypothesis generation, mechanistic insight, or validation. Similarity metrics are an essential component of data mining, and are used to identify relevant data. In computational biology, a wide variety of similarity metrics have been devised to maximize specificity and sensitivity in sequences alignment [1], proteomic mass spectrometry [2], evolutionary tree building [3], co-expression network creation [4], etc. These algorithms are typically used to facilitate comparing a data point against a curated library of experiments, which can lead to insight [5]. As modern data generation capabilities have created a deluge of potential data to compare against, exhaustive similarity search may become computationally prohibitive for inefficient algorithms. Therefore, efficient and accurate algorithms for computing similarity are necessary. For instance, the library of integrated network-based cellular signatures (LINCS) program has generated over one million gene expression experiments [6]. To compute the pairwise similarity between all experiments therefore requires 0.5 trillion similarity calculations.

When doing similarity-based computations on very large data, a significant drawback is that most of the pairwise comparisons yield a negative result, i.e. the two data points are not similar. An example of this is sequence alignment of genomic data. The current NCBI nr database contains *>* 78 million proteins (as of January 2017, release 80), the vast majority of which are not related to an input query. It would be a significant waste of time to perform the Smith-Waterman local alignment search against all 78 million sequences [7]. To overcome this limitation and enable large-scale data mining, sequence comparison algorithms commonly filter the set of sequences in the library prior to a more sensitive search. The BLAST algorithm requires a shared k-mer between the query sequence and candidate sequences from the library [8]. Only those proteins which contain a shared k-mer progress to a full alignment. This style of filtering candidates before a more computationally expensive scoring scheme is a common strategy which allows data mining to scale to ever-larger data volumes [9–11].

A second method to improve the speed of an algorithm is to adopt a faster core calculation. Most scientific data is stored as floating-point numbers; multiplication or division of floats is relatively slow. Therefore, optimizing an approach to minimize these will improve the computational speed. Bit operations (e.g. AND, OR, XOR) are dramatically faster than multiplication, yet require a restructuring of the basic approach or a data transform. The FastBit algorithm transforms data into bitmaps, then performs a hybrid compression to enable several common algorithmic operations (e.g. less than operator, histograms, exact pattern matching). This method is specifically designed to facilitate querying very large libraries with scientific data of high cardinalities [12]. Similarly, bit-vectors have been used to improve the speed of string matching [13].

We combine these two ideas in the Blazing Signature Filter (BSF), a new approach to prune unproductive pairwise similarity calculations and enable large-scale data mining. The BSF identifies signatures in digital data through bit representation (non-full precision) and bitwise operators. These binary operands are dramatically more efficient than floating-point multiplication and division in terms of CPU cycles per comparison. This simple heuristic allow us to remove > 98% of pairwise comparisons rapidly and therefore concentrate computational effort on pairs that are more likely to be meaningfully similar, enabling data mining tasks which previously appeared infeasible. We demonstrate the power of the BSF by computing the pairwise similarity of all publicly available LINCS datasets and identifying similarity of all genomes annotated by the Kyoto Encyclopedia of Genes and Genomes (KEGG).

## Results

Mining large data repositories to identify similar datasets is a common technical task. Depending on the number of comparisons to be done, the time involved in this simple task may be prohibitive. In most large-scale pairwise similarity searches (e.g., identifying similarity between all public transcriptomics datasets), the vast majority of pairs will be dissimilar. Thus, the most efficient way to speed such data mining explorations is to rapidly identify dissimilar pairs and remove them from the analysis pipeline. The purpose of the BSF is to identify pairwise similarity comparisons which are unlikely to be statistically meaningful. Our heuristic is to binarize the data and calculate a similarity metric on the binary data using bit operators, as bitwise computation is dramatically faster than floating point operations. In this way, the BSF can work as a front-end filter to computational analysis tools and dramatically speed up their pipeline.

### Algorithm description

A simplified example of the BSF is illustrated in Fig 1, where a 64 element signature is compared to a pool of candidates in a library. This 64 element signature is entirely binary, meaning that we only keep track of whether the element is part of the signature or not. Bit operations on two 64-bit binary signatures happen in a single instruction as two registers are compared with operators like AND, or XOR. Counting the number of successes in the comparisons (1s in the resulting array) is rapidly done using the hardware instruction ‘POPCNT’ [14]. In comparison, identifying the cosine similarity or Euclidean distance between two 64 element floating point vectors requires over 100 additions, multiplications, divisions and square root operations. For modern processors, cosine distance and euclidean distance have an average latency of 524*∼*538 and 711*∼*725 clock cycles, respectively, whereas BSF uses 4 clock cycles. Although this simplified example shows a 64 element signature, the software implementation of the BSF has been engineered to allow an arbitrary signature length. This is essential for comparing gene expression signatures which may scale to tens of thousands of elements, e.g. 20,000 human proteins. As this is larger than the size of a single CPU register, the data is chunked into appropriate sizes and comparisons are flowed through the registers.

**Fig 1.**
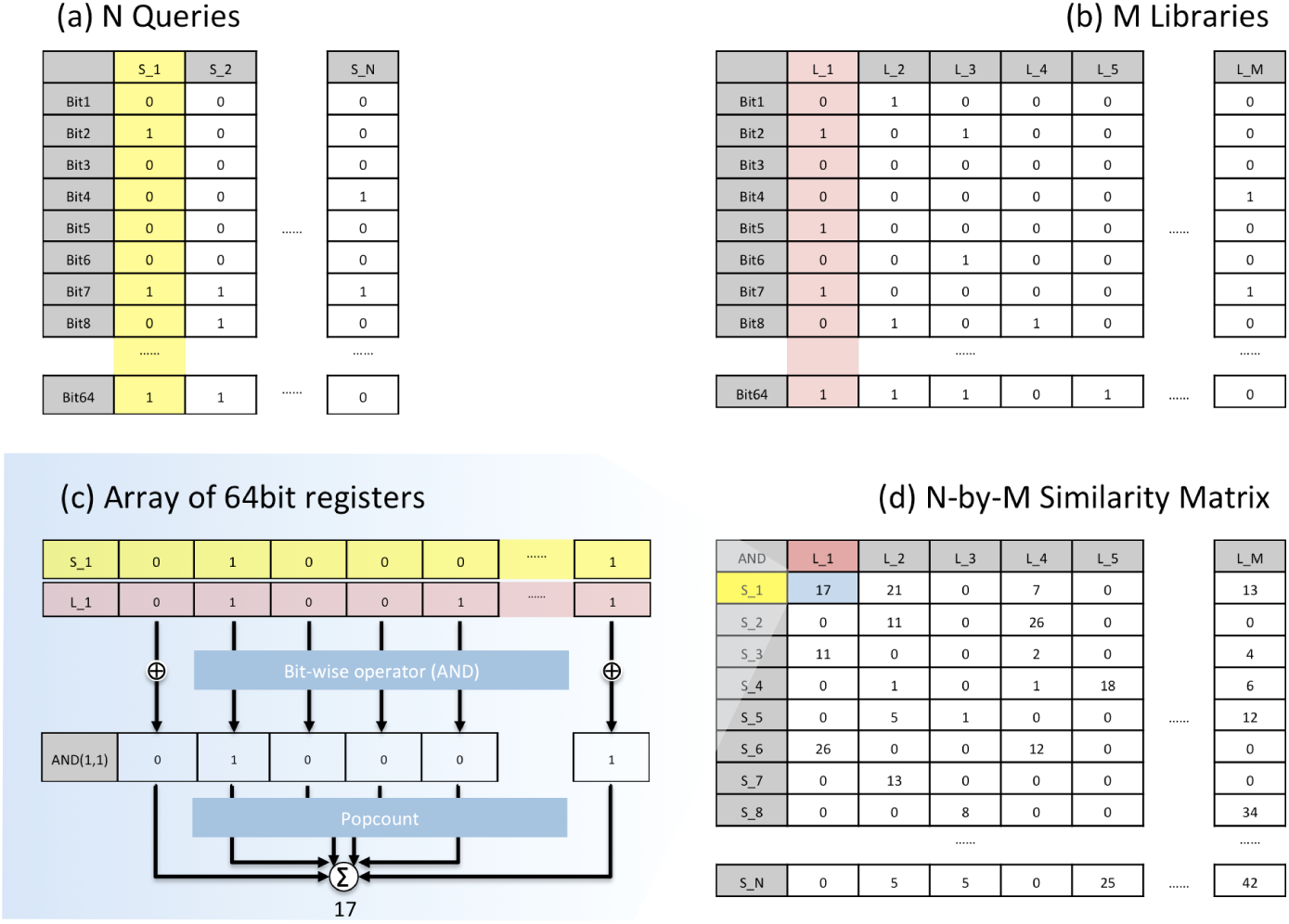
Illustration for the core of BSF. This simple example shows how the BSF deals with the binary data to identify the similarity between query and library signatures. Each signature has 64 elements (rows in the matrix). The binary data represents whether an element is part of the signature, i.e. ‘1’ means that the element is part of the signature, ‘0’ means that it is not part of the signature. (a) A set of query signatures represented as a binary matrix. (b) A set of library signatures to which the query signatures are compared. (c) The binary comparison for a single column in the query and library matrices. (d) The results matrix containing the similarity for each pairwise comparison. In the 64-bits example, clock cycle needs for the BSF are 1 for ‘AND’ and 3 for ‘POPCNT’, while cosine and euclidean metric use *>* 500 and *>* 700 clock cycles, respectively. (See Methods).

### Performance benchmarking

To demonstrate the speed of the similarity metrics, we performed a benchmark test of BSF, cosine similarity and Euclidean distance using a synthetic dataset mimicking gene expression measurements. In gene expression experiments, the goal is often to identify the up/down regulated genes relative to a reference condition. For example, in a gene knockout experiment, the desire is to understand and investigate which proteins are altered in their regulation relative to wild-type. The synthetic data was generated as measurements of 20,000 genes for 15,000 experiments (See Methods). Full precision floating point data was used by cosine and Euclidean distance metrics, whereas the BSF used binarized data showing up or down regulated genes.

We performed the full pairwise comparison of all 15K experiments for both the up and down matrix (*∼*225 million comparisons). To characterize the time-dependence of each algorithm on the length of the signature, we tested each algorithm with a different number of genes ranging from 1,000 to 20,000. This is essential to understanding the utility of each algorithm, as different applications may contain highly variable signature lengths. As shown in Fig 2, the time taken by each algorithm grows with the length of the signature. However, we note that the time dependence of the BSF grows dramatically more slowly than other methods. For the full 20,000 length signature (approximately what would be used for human gene expression data), the BSF algorithm ran in 45 seconds, while Euclidean and cosine methods took *∼*2,000 or *∼*6,000 seconds respectively. Both the Euclidean and cosine method show a linear time dependence on the signature length, *O*(*n*), while the BSF shows a log-linear dependence, *O*(log *n*), consistent with algorithmic expectations.

**Fig 2.**
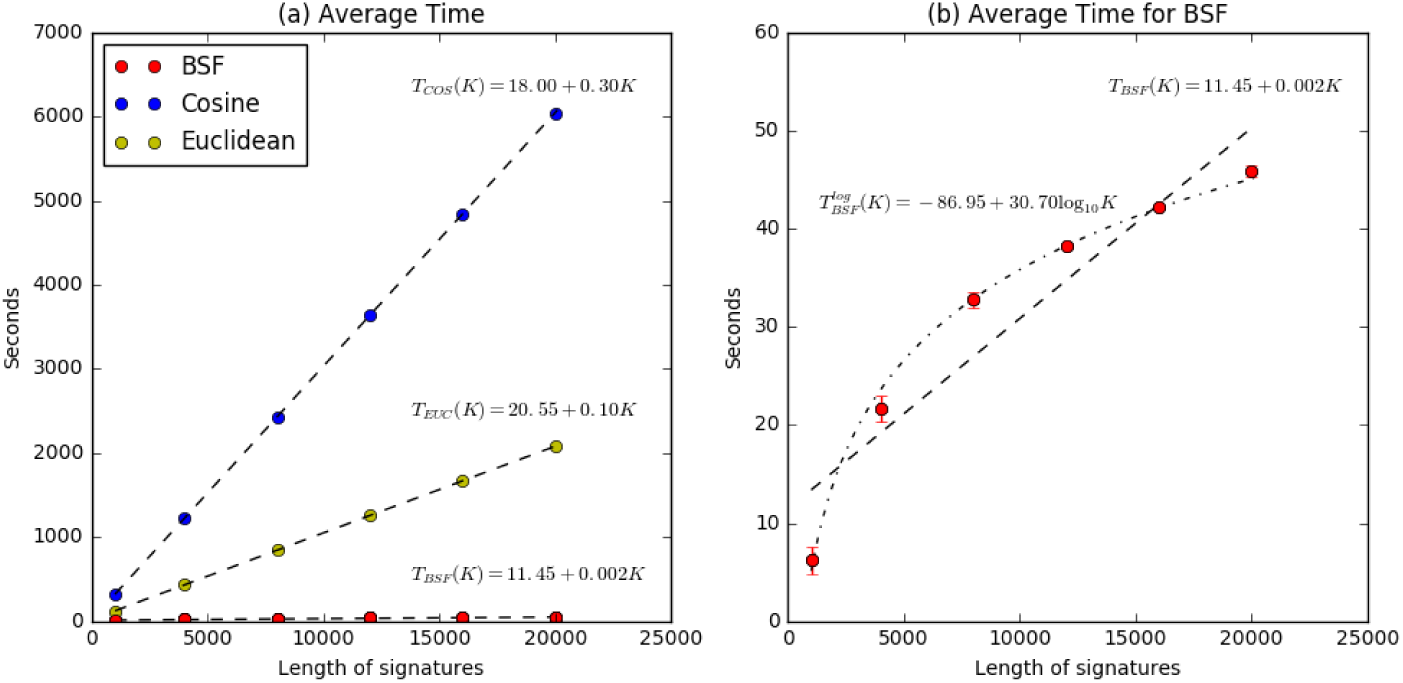
Benchmark result of BSF. (a) shows the average running time of computing each metric. *T*_*COS*_ (*K*), *T*_*EUC*_ (*K*), *T*_*BSF*_ (*K*) indicate the linear functions to fit the time points in terms of K of cosine similarity, euclidean distance, and BSF, respectively, where K means the length of each signature. (b) is a zoomed portion of (a) to focus on the BSF. (b) shows the BSF has a logarithmic time complexity while others have a linear time complexity. Refer to Methods and <u>https://github.com/PNNL-Comp-Mass-Spec/BSF_publication</u>.

### LINCS Network analysis

The LINCS L1000 project is a large-scale gene expression analysis, where numerous perturbations are applied to a variety of human cell lines and the response measured at multiple time points (<u>https://clue.io/</u>). The L1000 assay acquires transcript measurements on *∼* 1, 000 carefully chosen landmark genes followed by imputing the expression values for the remaining *∼* 21, 000 human genes. Perturbations used in the LINCS L1000 project include small molecule inhibitors, gene knockdowns and gene over-expression. Identifying similarities between perturbations is a primary focus of the project, and the goal is to enable the characterization of drug compounds having unknown targets as well as to identify signaling cross-talk and other gene expression changes induced by the perturbations.

As a real-world test for the BSF, we computed the pairwise similarity for the publicly available subset of the LINCS L1000 datasets [6]. We downloaded the December 2016 snapshot which contains 117K signatures as differentially expressed genes calculated using the characteristic direction method. We convert the up/down regulated genes into a 22,688-by-117,373 binary matrix and computed the 6.89 billion pairwise comparisons for the up-regulated genes and another 6.89 billion comparisons for the down-regulated genes. Results from these were merged to show the number of up/down regulated genes shared between two experiments. Supplementary Figure 1 shows that 98.8% of all pairs shared less than 10 up/down regulated genes. By spending about 2 hours determining this lack of pattern similarity using the BSF, accurate distances did not need to be calculated for these unproductive pairs, thus saving 9.6 days of computation.

After computing all pairwise comparisons within the LINCS dataset, we built a network connectivity graph to identify similar signatures of gene expression among the various perturbations. In exploring this graph, we first examined perturbations using small molecule histone deacetylase (HDAC) inhibitors. We queried the network using nine well-known HDAC inhibitors (belinostat, entinostat, mocetinostat, pracinostat, trichostatin A, vorinostat, rocilinostat, HDAC6 inhibitor ISOX, and valproic acid), which generated a sub-network of 1,066 nodes and 6.3 million edges. Fig 3 shows the network of the top 500 connections. Each node and its size indicate a perturbation dataset and the number of up/down-regulated genes by each perturbation. The weight of an edge shows the similarity score between two nodes. This analysis revealed that the nodes clustered by cell line rather than drug, indicating the response to various HDAC inhibitors is more cell line-specific than drug-specific. In addition to the query perturbations, six additional drug treatments were also found to show a similar signature and thus form part of the sub-network. Among these six are known or putative HDAC inhibitors such as HC toxin [15], panobinostat [16] and scriptaid, one of the first HDAC inhibitors discovered via high-throughput screening [17]. THM-I-94 had previously been hypothesized to act as an HDAC inhibitor based on structural similarity [18], and its clustering here supports that assertion. Other small molecules which cluster with the HDAC inhibitors include unnamed or poorly characterized pharmacological agents. With respect to the differential gene expression pattern shared by these drug treatments, we found enrichment in pathways associated with the cell cycle, MAPK signaling and KEGG’s Pathways in Cancer network based on the Fisher’s exact test.

**Fig 3.**
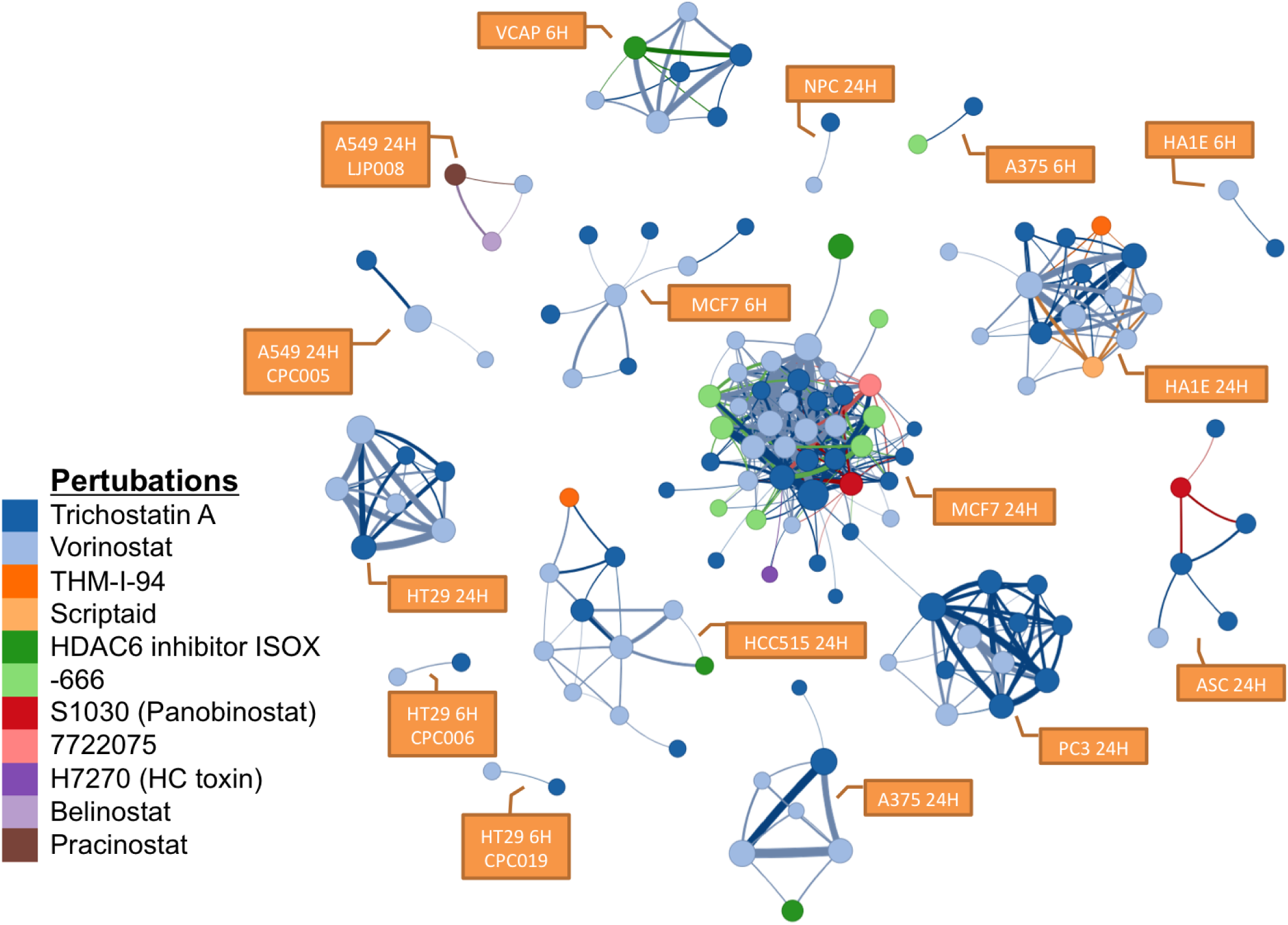
A sub-network associated to known HDAC inhibitors. The top 500 edges (among 6.3 million) are shown which includes the perturbations from the query (belinostat, pracinostat, trichostatin A, vorinostat, and HDAC6 inhibitor ISOX) and other compounds, some of which are known (scriptaid) and putative (THM-I-94) HDAC inhibitors. H7270 and S1030 are catalog numbers for HC toxin and panobinostat, both recognized HDAC inhibitors. Other perturbation are unnamed drugs (See Methods). The networks naturally form tight clusters, mostly distinguished by cell type and time point. The line width represents its similiarity score between two nodes.

A second data mining example from LINCS investigates the effect on human cell lines of non-human medication. Niclosamide is used to treat tapeworm infestations, but has recently been explored as a treatment for MRSA and Zika virus [19, 20]. With the capability of BSF, we can easily find the most similar treatments in the LINCS dataset. As shown in Supplementary Figure 2, 257 experiments of 20 different drugs form a subnetwork with niclosamide. Even though it wasn’t designed to target human cells, niclosamide has strong connectivity with IMD-0354, which is an IKK*β* inhibitor and blocks I*κ*B*α* phosphorylation. In addition to their high concordance in affecting the NF-*κ*B pathway (Supplementary Figure 2b), the two signatures have very high overlap in KEGG’s cell cycle pathway, with both showing strong down regulation of cyclins, cyclin dependent kinases, checkpoint kinases and the MCM complex (Supplementary Figure 2c).

### Whole genome similarity

A second application of the BSF is to compare gene content across a large number of genomes. Sequenced genomes are functionally annotated both by sequence repositories for inclusion in RefSeq [21] or Uniprot [22], or they can be annotated by a variety of systems biology style knowledgebases like KEGG [23] or RAST [24]. At the advent of genome sequencing, large scale comparisons of all genomes was used to understand protein function and evolution [25]. As genome sequencing technology has improved, the number of publicly available genomes grows dramatically and an all versus all comparison is much less frequently done due to computational costs.

KEGG is a functional annotation system which organizes whole genomes into into pathways and molecular interactions for 20,624 protein orthologs in 4,648 organisms (356 eukaryotes, 4,049 bacteria, and 243 archaea). Annotated genes are identified by their KEGG ortholog number, which are used to define metabolic, signaling and information pathways. Using the BSF, we computed the functional similarity between all genomes annotated by KEGG. Because gene presence/absence is already a binary measure, genome similarity comparisons are a simple and natural use for the BSF. We prepare the binary matrix consisting of 20,624 rows (orthologs) by 4,648 columns (genomes) where each cell represents where whether a ortholog is present in a genome. Computation for the full pairwise comparison took 5.2 seconds.

To show the diversity of genomic content within a taxonomic grouping, we plotted the average number of shared genes between genomes within a taxonomic group, e.g. Homo sapiens compared to all vertebrates (Fig 4). Eukaryotic genomes are generally larger than genomes of bacteria and archaea, and therefore it is not surprising to find a higher number of shared genes among eukaryotes. Additionally, KEGG contains a significant number of orthologs annotated in human disease pathways, and therefore the number of shared genes among animals is notably higher than among plants. We note that there is a broad range of similarity within a taxonomic group, most of which appears to be driven by genome size. For example, within alphaproteobacteria, most organisms share between 500-1100 orthologs. There are, however a few which share < 140 genes. These are 4 different strains of Candidatus Hodgkinia cicadicola (See Materials and Methods), an endosymbionts of the cicada, which has a tiny 144 kb genome [26]. To look at the comparisons within a taxonomic division, we plotted the average similarity between all genera within the class Bacilli (Fig 5). Many genera within Lactobacillales have very low similarity to all other genera of Bacilli. Some of this can be due to low gene counts (e.g. Weissella), however, several have high self-similarity but very low average overlap with any other taxa (e.g. Streptococcus, Leuconostoc, Melissococcus). Thus they likely represent an adaptive genomic response to unique environmental niche.

**Fig 4.**
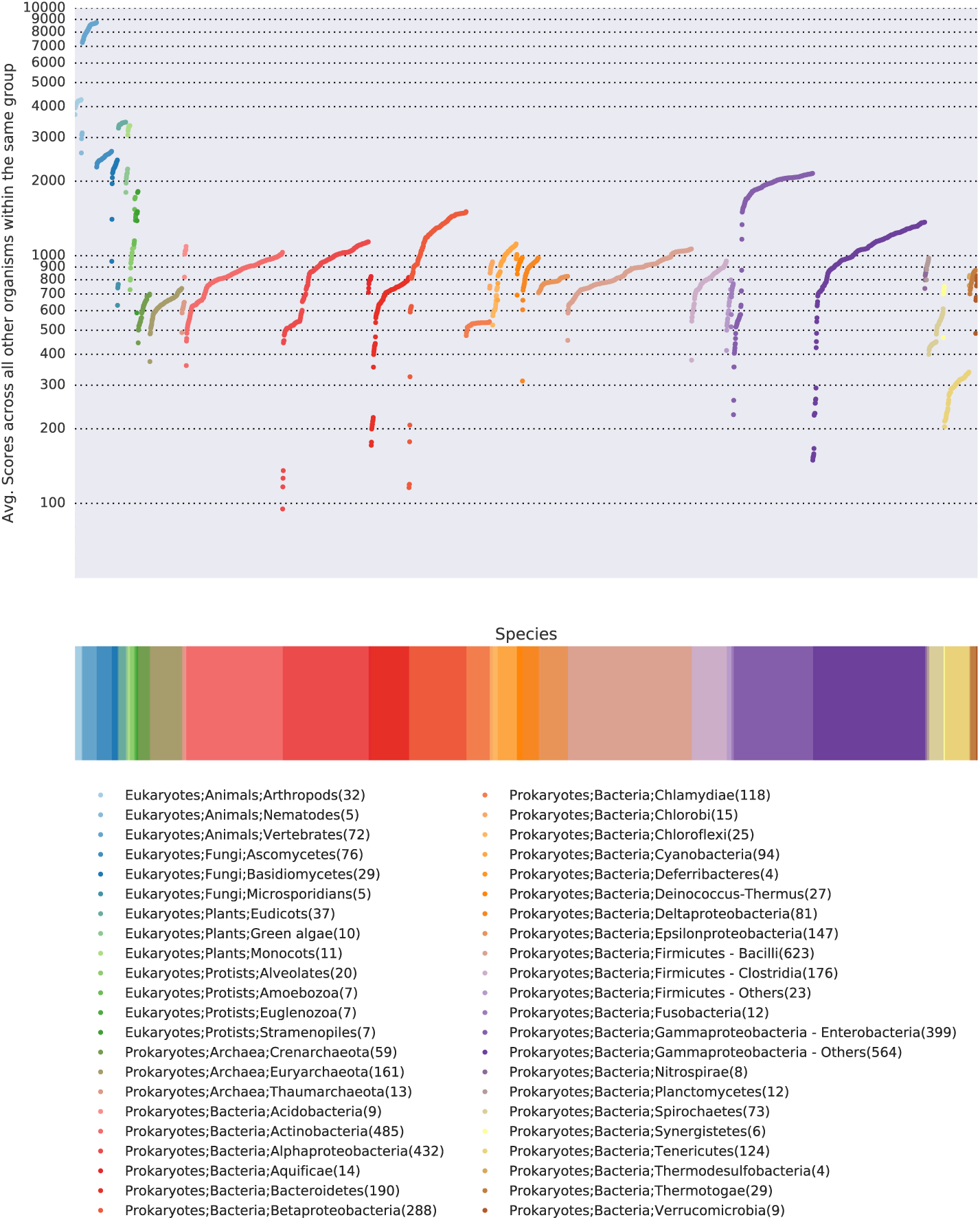
The average number of shared genes across all the genomes. Using protein orthologs from KEGG, we compare the gene presence/absence across all annotated genomes. Data points represent the average number of shared genes of a genome with the other genomes in the same taxonomic family. The graph is plotted on a log scale. Data points are colored according to taxonomic family membership. For clarity, taxonomic groups with fewer than four organisms were not plotted.

**Fig 5.**
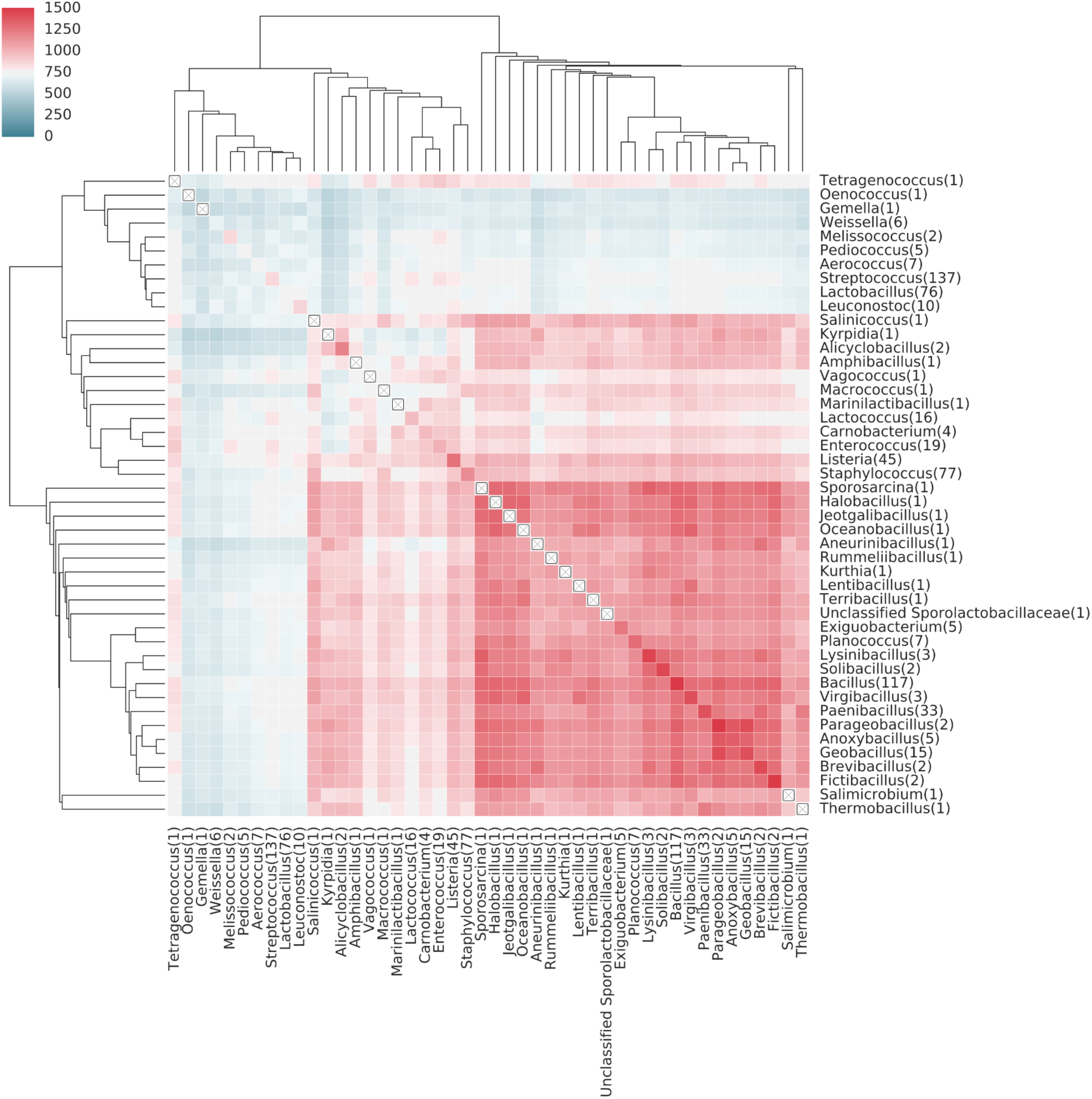
The average number of shared genes in genera within the class Bacilli. Rows and columns are ordered via hierarchical clustering. When only one sequenced member is present, the diagonal is marked with a boxed ×. A number in the bracket of each label shows the number of species of each genus. For details, see Materials and Methods.

Most organisms which share a small number of genes with other organisms are genome reduced and live as obligate symbionts. To compare the functions retained by genome reduced organisms, we plotted the similarity between organisms which had fewer than 600 genes annotated by KEGG (Fig 6). The lack of similarity between these minimalist genomes points to the wide variety of possible adaptations to environmental conditions. This is even true for parasites/pathogens which have nominally similar environments: e.g. Chlamydia and Mycoplasma both infect humans, Coxiella and Borrelia are both tick borne pathogens infecting humans.

**Fig 6.**
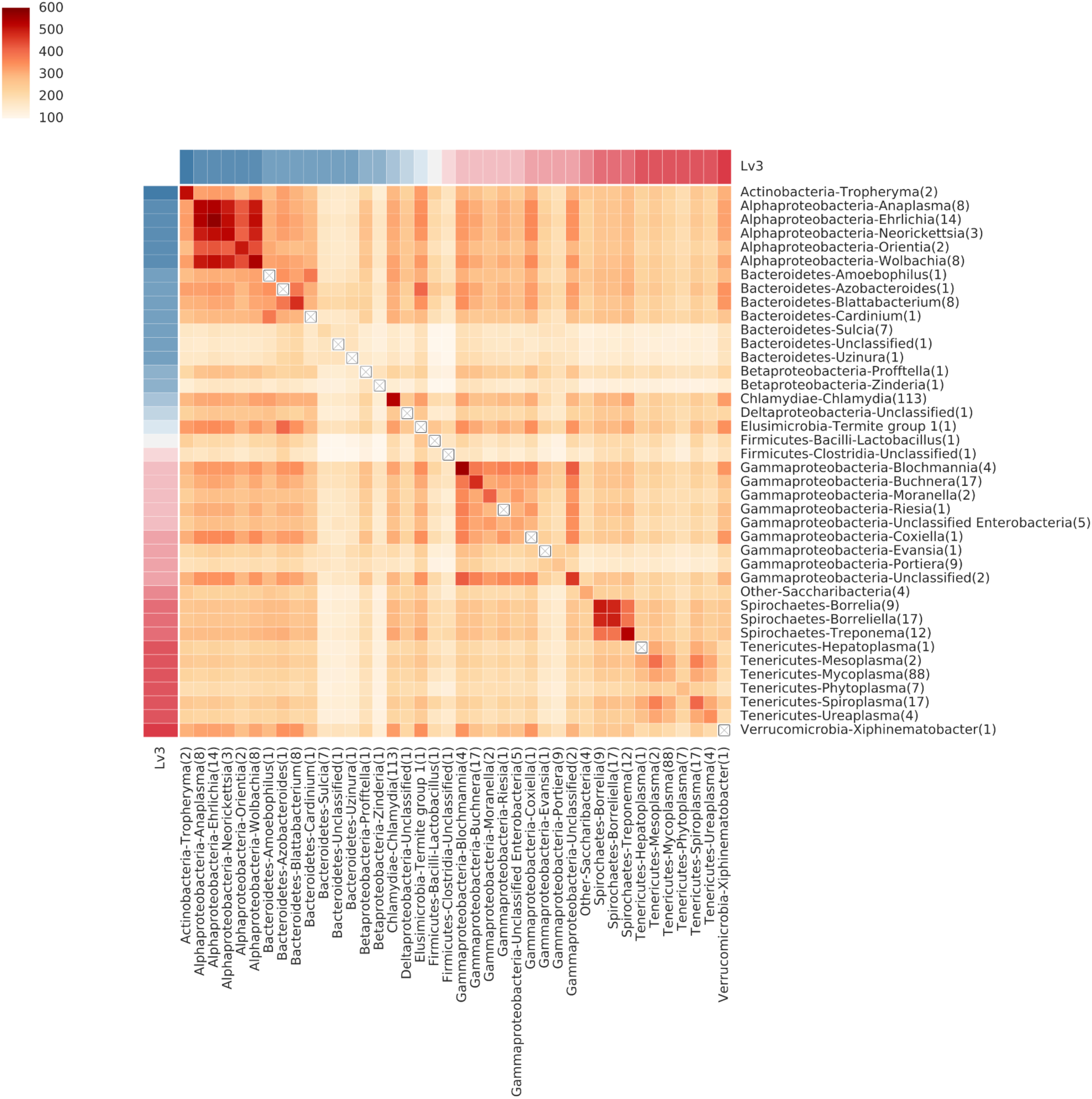
The average number of shared genes between minimalist bacteria. These organisms have 200-600 genes annotated by KEGG. Organisms are grouped by taxonomy.

## Discussion

As technologies for scientific data generation continue to dramatically improve and facilitate an ever greater characterization and description of the natural world, data mining for hypothesis generation and validation becomes both more important and more technically challenging. With the BSF, we introduce a simple and efficient method for identifying patterns, or signatures, in massive amounts of data. This is enabled by the rapid pairwise comparison of data as binary vectors. We show two example applications where pairwise comparisons are a common bioinformatics task: comparing genomes for similar gene content and identifying experiments with similar gene expression patterns. In both applications, the sheer number of comparisons would be time prohibitive without optimized computational methods such as the BSF.

New experimental technologies will improve the ability to make comprehensive datasets. For example, the task of identifying genetic interactions between pairs of genes was previously difficult to scale to whole genomes [27]. However, the CRISPR technology now makes is dramatically simpler to explore the effects of multiple knockouts [28], and we anticipate that whole genome double knockouts will be common in the near future. Even for genomically compact bacteria with ∼2000 genes, the number of double knockouts exceeds millions of strains. The subsequent task of identifying similarity (or differences) between the millions of strains will then require trillions of calculations. In these scenarios, efficient similarity metrics like the BSF will be essential to enable scientific discovery.

For datasets that are natively binary (e.g. gene content), the BSF works trivially. Another computation that is inherently binary is the set overlap calculation that is part of a Fisher’s exact test, commonly used for gene set enrichment. For datasets which are numeric or categorical, use of the BSF requires a meaningful transformation into binary space such as was done in the LINCS gene expression compendium. A wide variety of bioinformatics needs, e.g. proteomics library searches and FBA modeling, could benefit from using the BSF to quickly filter out unproductive data point prior to a more sensitive computation on the native (i.e. non-binary) data.

## Materials and Methods

The data and analysis methods for all figures are available at see https://github.com/PNNL-Comp-Mass-Spec/BSF_publication.

### LINCS Application

The LINCS L1000 project measures gene expression (transcriptomics) over different cell lines with a broad range of small molecule perturbations and genetic manipulations (knockout, knockdown and over-expression) [29, 30]. In this manuscript, we use the L1000 mRNA gene-expression signatures computed using the characteristic direction signatures method [6, 29], giving binary up and down regulated genes for each of the ∼ 117, 000 datasets. It is publicly available at <u>http://amp.pharm.mssm.edu/public/L1000CDS_download/</u>.

### KEGG Application

All Kegg annotations were taken from in Release 81.0 downloaded on January 1, 2017. A table of orthologs versus genomes was created and fed into the BSF using the python interface. See the kegg data section at https://github.com/PNNL-Comp-Mass-Spec/BSF_publication for auxiliary files and code.

### Benchmarking

The synthetic benchmarking data was created as a table (15*K* columns × 20*K* rows) of floating point numbers drawn randomly from the gaussian distribution of *N* (0, 0.5). Rows can be thought of as different gene measurements, and columns as distinct datasets. This continuous data was binarized into two tables to represent the extremes of the distribution, i.e. values < *-*0.6 were written as 1 in a binary table representing the ‘low’ values and values > 0.6 were written as 1 to a binary table representing high values.

We use the cosine distance and euclidean distance on the original floating point data to compare the performance with BSF. In the manuscript, we discussed clock cycles required for various operations. For a description of CPU clock cycles per instruction set, refer to http://www.agner.org/optimize/instruction_tables.pdf. Supposing the *M*-by-*N* signature matrix [*S*_1_, *S*_2_, *S*_3_, ··, *S*_*N*_], the formulae for cosine and Euclidean similarity are:

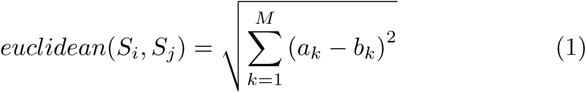

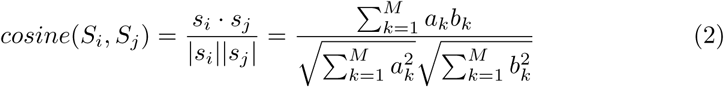

, where *S*_*i*_ = [*a*_1_, *a*_2_, *a*_3_,*…, a*_*M*_]^*T*^ and *S*_*j*_ = [*b*_1_, *b*_2_, *b*_3_, *…, b*_*M*_]^*T*^ (*i, j* = 1, 2, 3, *…, N*).

### BSF Software and Access

The BSF code, written in C++, is an open source software project licensed under BSD. Source can be found at https://github.com/PNNL-Comp-Mass-Spec/bsf-core. For ease of access, we have written a python wrapper to interface with the BSF C++ library, <u>https://github.com/PNNL-Comp-Mass-Spec/bsf-py</u>. Python extensions and numpy C-API are employed to implement the python wrapper.

The BSF is accessed through an API which ensures that input data is appropriate and meaningful, and interprets the output tables which are returned. Input to the BSF consists of two binary tables of size *K*-by-*N* and *K*-by-*M*. For the input table, all the bits of each column are stored into an array of 64-bit unsigned integers. It enables bitwise operators using the 64-bit registers. The user also specifies which binary operator to use. The return from BSF is an *N*-by-*M* table with each cell representing the value of comparing element *i* in table 1 with element *j* in table 2.

As shown in Fig 1, the BSF outputs the *N*-by-*M* matrix, where *N* and *M* indicate the column sizes of *K*-by-*N* and *K*-by-*M* input matrices, respectively. The pseudocode is described in Algorithms 1.

In case to compute all the pairwise comparisons between two signatures within a *K*-by-*N* library matrix, it outputs a *N*-by-*N* strictly upper triangular matrix with zero diagonal entries so that we avoid the redundant computation of *a*_*ij*_ for *i* > *j* which must be equal to *a*_*ji*_. The pseudo code for this looks as shown in Algorithms 2.

In case of LINCS dataset, we need 117*K* × 117*K* × 4bytes ≈ 50*G* at least to save all the results. As a data size gets larger, this may lead to an out-of-memory exception depending on the physical memory size. To safely avoid this memory issue, we split the output matrix into multiple chunks, of which size is reasonably manageable. Given the file size, the size of a chunk matrix is decided. Details are described in Algorithms 3.

#### Algorithm 1 BSF algorithm to analyse the similarity between columns of a library matrix and a query matrix

~~~
**Input:** *lib*[*N*_*row*_][*N*_*col*_] *←* a libary matrix, *q*[*N*_*row*_][*M*_*col*_] *←* a query matrix
**Output:** *out*[*N*_*col*_][*M*_*col*_] *←* a 2D array of unsigned integers
1: **function** BSFCORE1(*lib, q*)
2: **for all** *i* in [0, *N*_*col*_] **do**
3:     **for all** *j* in [0, *M*_*col*_] **do**
4:         *popcount ←* 0
5:         **for all** *k* in [0, *N*_*row*_] **do**
6:           *sim ← lib*[*k*][*i*] & *q*[*k*][*j*]
7:           *pop*_*count*_ *← pop*_*count*_ +̲̲builtin ̲popcountll(*sim*)
8:       *out*[*i*][*j*] *← count*
9: **return** out
~~~

#### Algorithm 2 BSF algorithm to analyse the similarity between columns of a single library matrix

~~~
**Input:** *lib*[*N*_*row*_][*N*_*col*_] *←* a 2D array of unsigned 64bit integers
**Output:** *out*[*N*_*col*_][*N*_*col*_] *←* a 2D array of unsigned integers
1: **function** BSFCORE2(*lib*)
2: **for all** *i* in [0, *N*_*col*_ *-* 1] **do**
3:    **for all** *j* in [*i* + 1, *N*_*col*_] **do**
4:       *popcount ←* 0
5:       **for all** *k* in [0, *N*_*row*_] **do**
6:         *sim ← lib*[*k*][*i*] & *lib*[*k*][*j*]
7:         *pop*_*count*_ *← pop*_*count*_ +̲̲builtin ̲popcountll(*sim*)
8:       *out*[*i*][*j*] *← count*
9:    **return** out
~~~

#### Algorithm 3 Chunk split for avoiding out-of-memory in handling a big library matrix

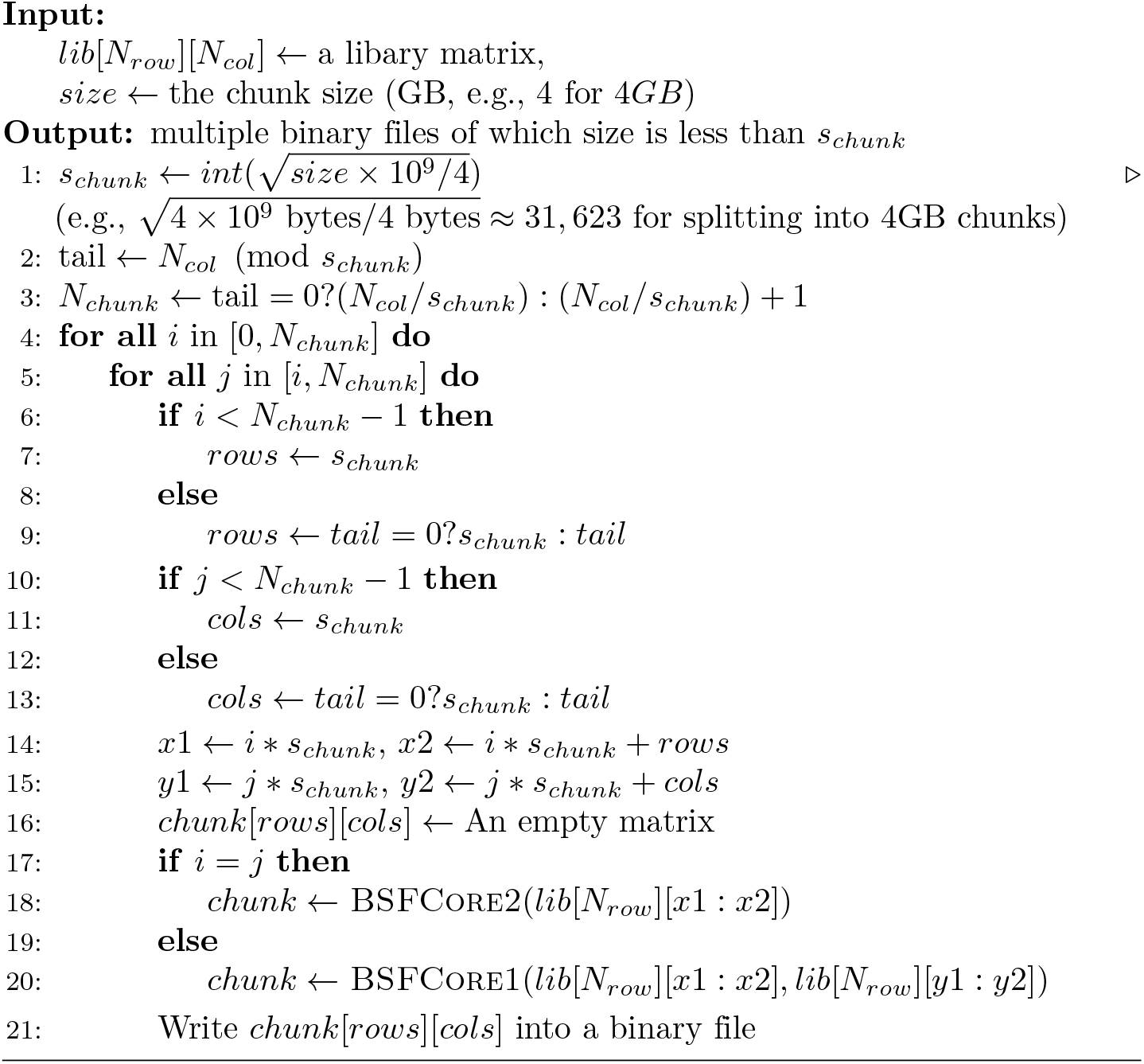

## Supporting information

**Supplementary Figure 1.**
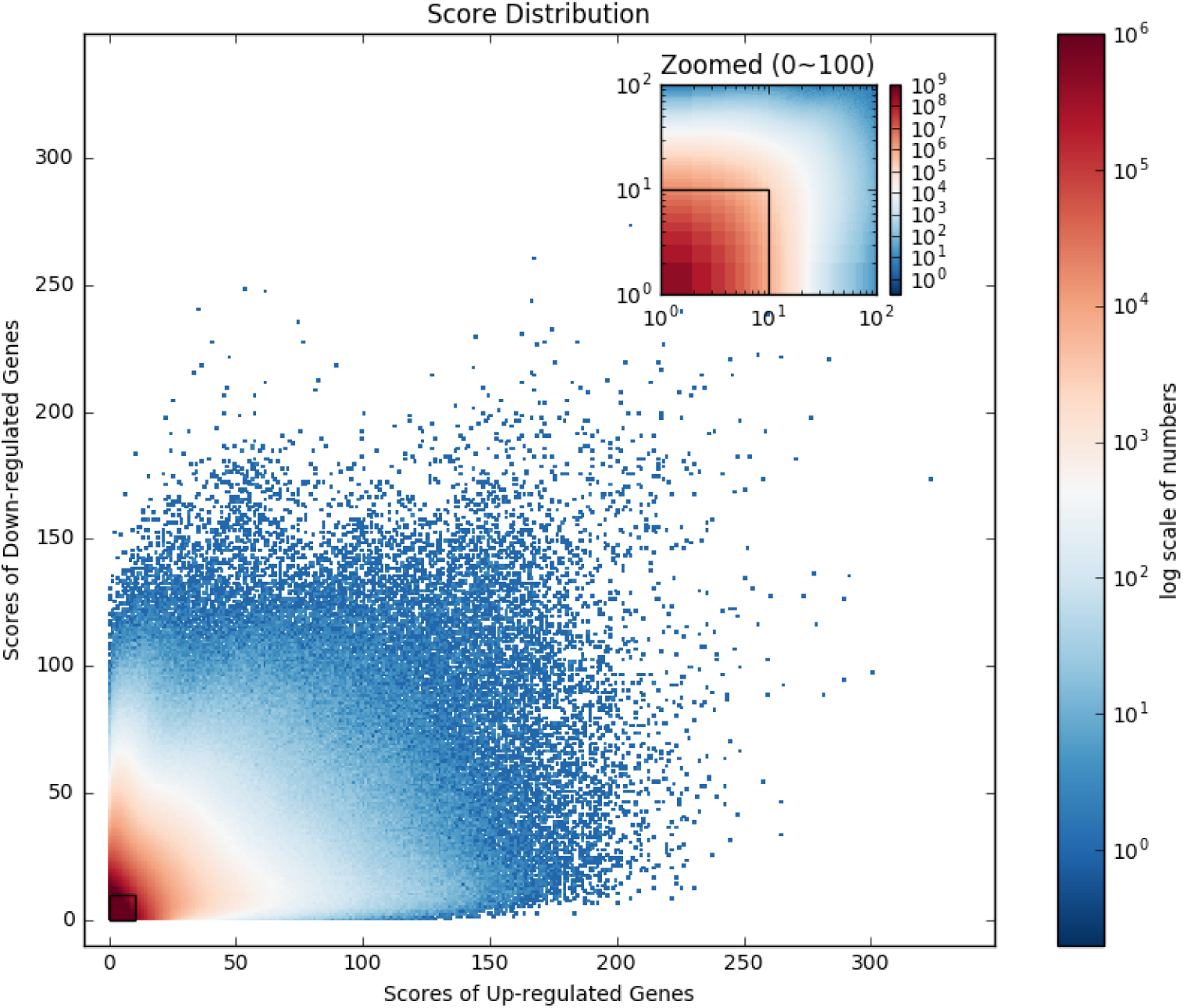
Score distribution of the 6.89 billion pairwise comparisons in the LINCS L1000 dataset. The color of each point describes the number of pairs which have shared genes. X and Y axes indicate the number of shared up-regulated genes and down-regulated genes, respectively. For example, a point of (50, 50) has 147, which means 147 pairs of two signatures share 50 up-regulated genes and 50 down-regulated genes. The overwhelming majority of pairwise comparisons, ∼6.80 billions or 98.8%, are located in a small box of up-regulated genes < 10 and down-regulated genes < 10. These represent pairs of experiments, which do not share a discernable signature of regulated gene expression and are unproductive data mining events.

**Supplementary Figure 2.**
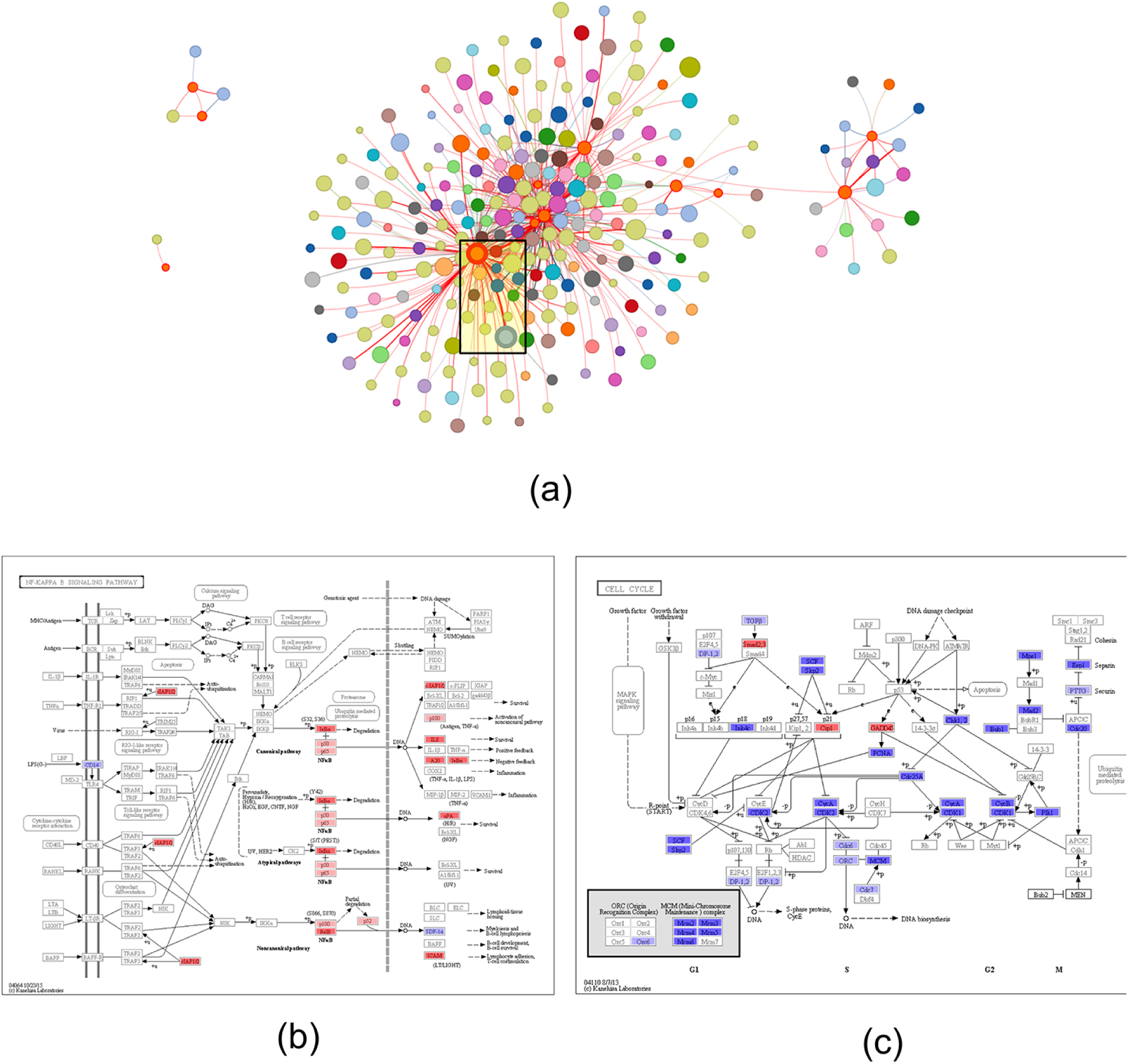
A sub-network of LINCS L1000 experiments most similar to niclosamide. (a) We extracted the network for 88 datasets associated with non-human medications such as niclosamide (tapeworm infestations) and daminozide (plant growth regulator). It shows 257 experiments of 20 drugs highly connected to these 88 signatures. Refer to Materials and Methods for details. (b) Differentially expressed genes shared between niclosamide and IMD 0354, an IKK*β* inhibitor. Most of all common genes are down-regulated and cell cycle looks slow down. (c) Shared differential genes shown for the NF-*κ*B signaling pathway; most of the genes are up-regulated.

## Acknowledgements

This work was supported by the National Cancer Institute (NCI) CPTAC award U24 CA210972 (SHP), and the U.S. Department of Energy, Office of Science, Office of Biological and Environmental Research, Pan-omics Program. Battelle operates the Pacific Northwest National Laboratory for the DOE under contract DE-AC05-76RLO01830.

